# Butyryl/Caproyl-CoA:Acetate CoA-Transferase: Cloning, Expression and Characterization of the Key Enzyme Involved in Medium-Chain Fatty Acid Biosynthesis

**DOI:** 10.1101/2021.03.10.434813

**Authors:** Qingzhuoma Yang, Shengtao Guo, Qi Lu, Yong Tao, Decong Zheng, Qinmao Zhou, Jun Liu

**Affiliations:** Key Laboratory of Environmental and Applied Microbiology, Environmental Microbiology Key Laboratory of Sichuan Province, Chengdu Institute of Biology, Chinese Academy of Science, Chengdu 610041, China; University of Chinese Academy of Sciences, Beijing 100049, China; BGI Education Center, University of Chinese Academy of Sciences, Shenzhen 518083, China; Faculty of Bioengineering, Sichuan University of Science & Engineering, Xueyuan Street 180^#^, Huixing Rd. 643000, Zigong, P.R. China

**Keywords:** CoA-transferase, chain elongation, Medium-chain Fatty Acids, *Ruminococcaceae* bacterium, Caproic acid

## Abstract

Coenzyme A transferases (CoATs) are important enzymes involved in carbon chain elongation contributing to medium-chain fatty acid (MCFA) biosynthesis. For example, butyryl-CoA:acetate CoA transferase (BCoAT) is responsible for the final step of butyrate synthesis from butyryl-CoA. However, little is known about caproyl-CoA:acetate CoA-transferase (CCoAT), which is responsible for the final step of caproate synthesis from caproyl-CoA. In this study, two CoAT genes from *Ruminococcaceae* bacterium CPB6 and *Clostridium tyrobutyricum* BEY8 were identified by gene cloning and expression analysis. The enzyme assays and kinetic studies were carried out using butyryl-CoA or caproyl-CoA as the substrate. CPB6-CoAT can catalyze the conversion of both butyryl-CoA to butyrate and caproyl-CoA to caproate, but its catalytic efficiency with caproyl-CoA as the substrate was 3.8 times higher than that with butyryl-CoA. In contrast, BEY8-CoAT had only BCoAT activity, not CCoAT activity. This demonstrated the existence of a specific CCoAT involved in chain elongation via the reverse β-oxidation pathway. Comparative bioinformatics analysis showed the presence of a highly conserved motif (GGQXDFXXGAXX) in CoATs, which is predicted to be the active center of CoATs. Single point mutations in the conserved motif of CPB6-CoAT (Asp346 and Ala351) led to marked decreases in the activity for butyryl-CoA and caproyl-CoA, indicating that the conserved motif is the active center of CPB6-CoAT, and sites Asp346 and Ala351 were critical residues that affect enzymatic activity. This work provides insight into the function of CCoAT in caproic acid biosynthesis and improves the understanding of the chain elongation pathway for MCFA production.

## Introduction

Medium-chain fatty acids (MCFAs, C6-C12) are widely utilized in agriculture and industry. For example, *n*-caproic acid (C6) is used as a precursor for the production of fragrances (1), antimicrobial agents (2), and drop-in biofuels (3). Recent studies have shown that MCFAs produced from renewable feedstock by anaerobic fermentation hold promise for replacing fossil resources and botanical oils such as palm kernel oil to meet the requirements for sustainable development (4). A few microorganisms, such as *Megasphaera elsdenii* (5), *Ruminococcaceae* bacterium CPB6 (6), *Acinetobacter* spp. (7), and *Clostridium kluyveri* (8), have been reported to be able to synthesize MCFAs from renewable feedstock via the carbon chain elongation pathway (9). In the process of chain elongation, intermediates of acidogenesis, such as acetate (C2) and *n*-butyrate (C4), as substrates are elongated to caproic acid (C6) and octanoic acid (C8) by adding acetyl-CoA in reverse β-oxidation cycles (10,11). C2 or C4, transformed to acetyl-CoA or butyryl-CoA, respectively, represents the initial substrate for elongation in reverse β-oxidation. The pathway has been identified as a key metabolic process in MCFA biosynthesis (12).

The production of high concentrations of butyrate (>10 mM) in vitro has been reported in some anaerobes, such as *Roseburia* (13) and *Faecalibacterium* (14). Butyrate is normally generated from two molecules of acetyl-CoA, yielding acetoacetyl-CoA, which is then converted to butyryl-CoA (15). In the latter reaction, butyryl-CoA is exchanged with exogenously derived acetate to yield acetyl-CoA and butyrate (16). The enzymes responsible for butyrate production in the reverse β-oxidation pathway comprise acetyl-CoA acetyltransferase (AtoB), 3-hydroxybutyryl-CoA dehydrogenase (Hbd), enoyl-CoA hydratase (Crt), butyryl-CoA dehydrogenase (Bcd), and butyryl-CoA:acetate CoA-transferase (BCoAT) (17). Among them, BCoAT is a well-known CoA-transferase (CoAT) responsible for the final step of butyric acid synthesis, transforming the CoA moiety from butyryl-CoA to an exogenous acetate molecule, which results in the formation of butyrate and acetyl-CoA (18,19). CoATs are abundant in anaerobic fermenting bacteria that cope with low ATP yields, but they are also found in aerobic bacteria and in the mitochondria of humans and other mammals (20). The synthesis pathway and key genes associated with butyric acid in MCFA biosynthesis via reverse β-oxidation are well understood. However, little is known about key genes involved in the conversion of butyric acid (C4) to caproic acid (C6). Although most genes responsible for butyric acid production are suggested to function in further chain elongation of MCFAs (17), the fact that many butyrate-producing bacteria, such as *Clostridium tyrobutyricum*, produce only butyric acid instead of caproic acid via the reverse β-oxidation pathway suggests that there may be different functional genes involved in the production of caproic acid.

Recently, our study showed that *Ruminococcaceae* bacterium CPB6 is a caproic acid-producing bacterium with the highly prolific ability to perform chain elongation and can produce caproic acid (C6) from lactate (as an electron donor) with C2-C4 carboxylic acids and heptoic acid (C7) with C3-C5 carboxylic acids as electron acceptors (EAs) (21,22). Moreover, a set of genes correlated with chain elongation were identified by sequencing and annotating the whole genome of the CPB6 strain (23). However, very little information is available on enzymes involved in the conversion of C4 to C6, especially the gene responsible for the conversion of caproyl-CoA to caproic acid.

In this study, we cloned a predicted CCoAT gene from the caproic acid-producing strain CPB6 (21) and a BCoAT gene from the butyric acid-producing *C. tyrobutyricum* BEY8 (24) and expressed the two proteins in *Escherichia coli* BL21 (DE3) with the plasmid pET28a. The aims of this study were to (i) compare differences in sequence, structure, enzymatic activity and substrate specificity between the CCoAT and BCoAT; (ii) identify the active center of the CCoAT and its effects on the activities of enzymes with different structures; and (iii) verify the existence of the CCoAT in the caproic acid biosynthesis pathway.

## Results and Discussion

### Cloning, expression, and purification of CoA-transferase

According to the genome sequences of strains CPB6 and *C. tyrobutyricum* BEY8, specific primers targeting CoAT genes were designed and synthesized (Table 1). Agarose gel electrophoresis showed that the size of the PCR products and the double-digestion products was approximately 1300 bp, consistent with the expected sizes of CPB6-CoAT (1344 bp) (Fig. S1) and the BEY8-CoAT gene (1233 bp) (Fig. S2). Sequence analysis of the recombinant CoAT plasmids showed that the cloned genes shared 100% similarity with the predicted CoAT genes of strains CPB6 (CCoAT) and BEY8 (BCoAT). This finding indicated that the recombinant *E. coli*/pET28a-CCoAT and *E. coli*/pET28a-BCoAT were successfully constructed.

**Table 1.**
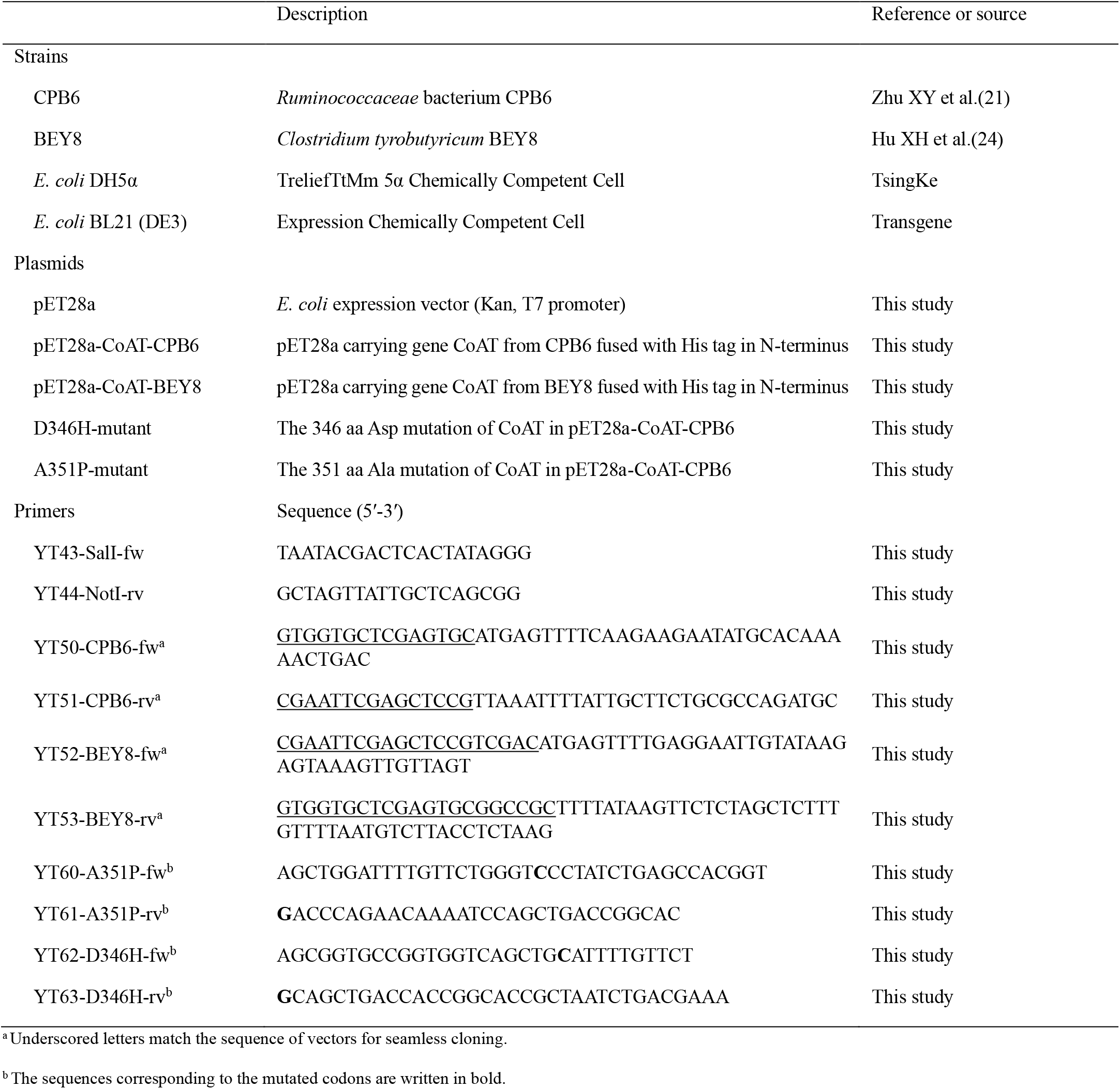
Bacterial strains, plasmids, and primers used in this study.

To characterize the functions of CoAT proteins, the two recombinant plasmids (pET28a-CCoAT and pET28a-BCoAT) were expressed in *E. coli* BL21 (DE3). Single bands of the purified proteins were detected on SDS-polyacrylamide gels after affinity chromatography (Fig. 1*A*). As shown in Fig. 1*A*, there was no obvious protein band of approximately the size of the target protein in *E. coli*/pET28a (control), while a single band was observed in *E. coli*/pET28a-BCoAT (lane 2) and *E. coli*/pET28a-CCoAT (lane 3), and their sizes were consistent with the expected sizes of BEY8-CoAT (46 kDa) and CPB6-CoAT (49 kDa). Furthermore, western blotting analysis with a His-antibody (Fig. 1*B*) also demonstrated that the observed bands were consistent with the expected molecular mass of BEY8-CoAT and CPB6-CoAT (approximately 46-49 kDa).

**FIGURE 1.**
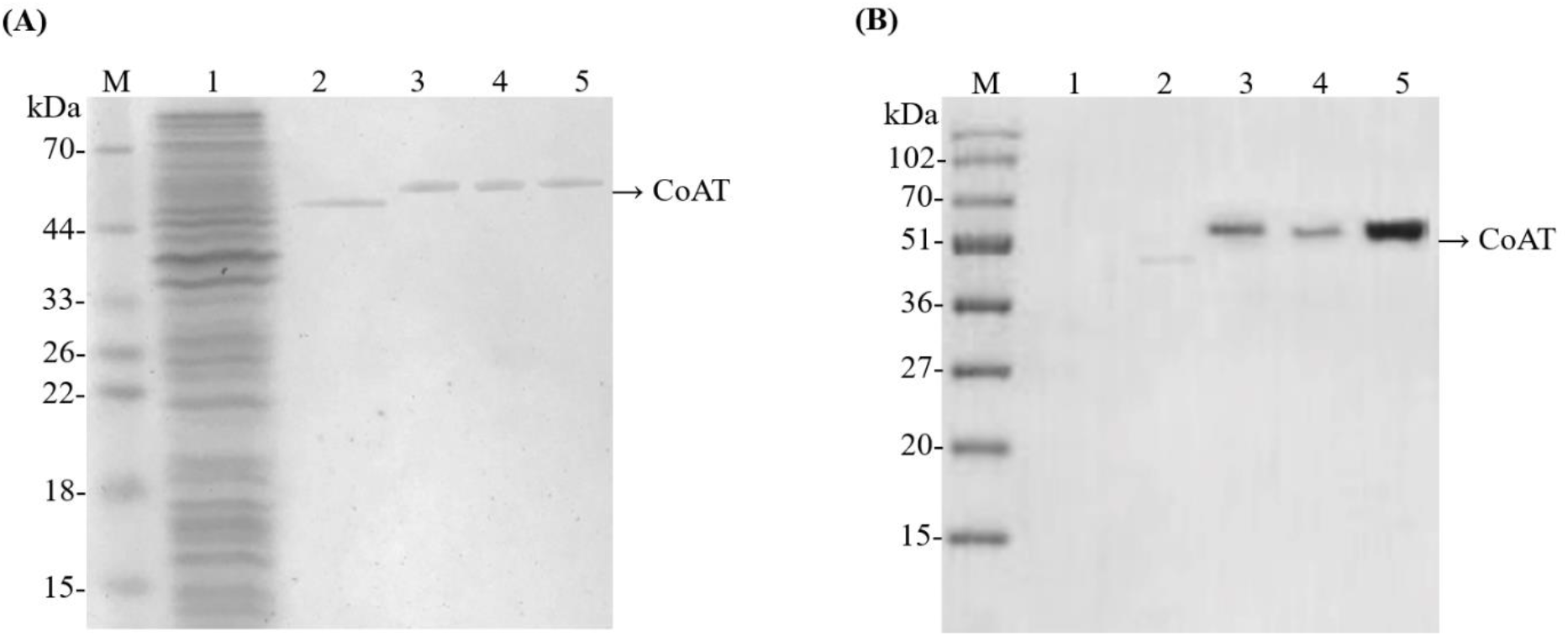
Purification and western blot analysis of CPB6-CoAT (CCoAT) and BEY8-CoAT (BCoAT). Analysis of the purified CCoAT and BCoAT by SDS-PAGE **(A)**. Analysis of the purified CCoAT and BCoAT by western blotting with anti-His-tag antibody **(B)**. M, molecular mass marker. Lanes: 1, pET28a; 2, BCoAT; 3, CCoAT; 4, CCoAT-D346H mutant; 5, CCoAT-A351P mutant. Samples (∼2 µg) were visualized by Coomassie Brilliant Blue staining after electrophoresis. Molecular mass positions are shown by markers (kDa).

### Enzyme assay

The CoAT activity of crude enzyme extracts was determined by measuring the production of acetyl-CoA from butyryl-CoA or caproyl-CoA (25). It has been previously reported that the key reactions for butyrate and caproate production were (1) butyryl-CoA + acetate→ butyrate + acetyl-CoA and (2) caproyl-CoA + acetate → caproate + acetyl-CoA (10,21,26). As shown in Table 2, the crude and purified BEY8-CoAT activities with butyryl-CoA and sodium acetate as substrates were 6.91±0.12 and 26.2±0.09 U/mg of protein, respectively. However, this enzyme showed no activity for caproyl-CoA. This result suggested that BEY8-CoAT is a BCoAT, similar to the CoAT from *Clostridium acetobutylicum* ATCC 824 that is able to produce butyrate instead of caproate, and its purified enzyme activity was 29.1 U/mg of protein (27). Moreover, the butyrate-producing bacterium *Coprococcus* sp. strain L2-50 from the human large intestine showed very high BCoAT activity (118.39±5.02 U/mg of protein) but no CCoAT activity (13). This result indicated that the BCoAT probably has substrate specificity for butyryl-CoA (29). In contrast, the activities of crude and purified CPB6-CoAT with butyryl-CoA and sodium acetate as substrates were 2.07±0.06 and 10.8 ±0.02 U/mg of protein, and the activities with caproyl-CoA and sodium acetate as substrates were 5.11±0.08 and 27.6±0.15 U/mg of protein, respectively (Table 2), indicating that CPB6-CoAT can catalyze the conversion of both butyryl-CoA to butyrate and caproyl-CoA to caproate. It is worth noting that the crude and purified CPB6-CoAT activity for caproyl-CoA was 2.5-2.6 times higher (5.11 vs 2.07, 27.56 vs 10.28 U/mg of protein) than that for butyryl-CoA, suggesting that CPB6-CoAT specifically prefers caproyl-CoA as a substrate instead of butyryl-CoA.

**Table 2.** Purification and specific activities of the CoATs^*a*^.

| Enzyme | Total protein, mg | Total activity, U | Specific activity, U/mg of protein |  | Purification fold |
| --- | --- | --- | --- | --- | --- |
|  |  |  | Butyryl-CoA | Caproyl-CoA |  |
| The CPB6-CoAT |  |  |  |  |  |
| Crude extract | 87.04 | 180.2 | 2.07±0.06 | 5.11±0.08 | 1 |
| Purified protein<sup>b</sup> | 22.61 | 244.2 | 10.8±0.02 | 27.6±0.15 | 5.4 |
| The BEY8-CoAT |  |  |  |  |  |
| Crude extract | 63.10 | 436.0 | 6.91±0.12 | ND<sup>a</sup> | 1 |
| Purified protein<sup>b</sup> | 17.66 | 462.7 | 26.2±0.09 | ND<sup>a</sup> | 3.8 |

<sup>a</sup>The purification data in the table were from 300 ml of culture medium. U = μmol/min. ND = not detectable.
<sup>b</sup>Protein was purified with affinity chromatography.

The BCoAT is required for butyrate biosynthesis in *Clostridium kluyveri* (13) and *C. tyrobutyricum* (28). This enzyme is responsible for the final step of butyrate production, catalyzing the conversion of butyryl-CoA and acetate to butyrate and releasing acetyl-CoA (16). As reported in previous studies, this enzyme is considered to be a biomarker for identifying butyrate-producing bacteria (16,27,29). However, BCoAT is not responsible for chain elongation of larger or higher-carbon-numbered (>C5) fatty acids (28). Seedorf et al.(17) speculated that BCoAT may catalyze the conversion of caproyl-CoA to caproate, similar to the conversion of butyryl-CoA to butyrate, based on genome analysis of *C. kluyveri*, but no further research has been reported. Our previous study showed that the rate of caproate production with caproyl-CoA as the substrate in strain CPB6 was 3.5 times higher than that observed with butyryl-CoA as the substrate and suggested the existence of a CCoAT that specifically prefers caproyl-CoA instead of butyryl-CoA as the substrate (21). In this study, CPB6-CoAT was confirmed for the first time to be a CCoAT responsible for the final step of caproate formation, although it harbored low BCoAT activity for butyryl-CoA. These data demonstrated the existence of a specific CCoAT involved in the chain elongation of MCFAs, which is significantly different from the function of BCoAT. The detailed mechanism underlying this functional difference needs to be further studied.

### Kinetics of CoA-transferases

The kinetic parameters of the recombinant proteins were investigated using a colorimetric assay according to a previous study (16). Initial velocities were determined at fixed sodium acetate concentrations with different butyryl-CoA or caproyl-CoA concentrations. *K*_m_ and *V*_m_ values were estimated from secondary plots (Materials and Methods). Additionally, *k*_cat_ values were calculated from enzyme concentrations in the reaction mixtures. The double-reciprocal plotting of enzyme kinetics showed that the reactions of the two CoATs follow the ternary-complex mechanism (Fig. S3), the result suggested that the CoATs of the CBP6 and BEY8 probably belong to family III, which can be distinguished kinetically(18).

As *k*_cat_/*K*_m_ can be used to compare the catalytic efficiency of different substrates catalyzed by the same enzyme (30), a lower *K*_m_ value indicates that the enzyme has a higher affinity for the substrate and vice versa (31). In this study, the Km, *k*_cat_, *k*_cat_/*K*_m_ and *V*_m_ values of CPB6-CoAT with caproyl-CoA were 358.8 µM, 14.74 min^−1^, 41.08 mM^−1^min^−1^ and 29.51 μM min^−1^, respectively, and those with butyryl-CoA were 536.9 µM, 5.810 min^−1^, 10.82 mM^−1^min^−1^ and 11.62 μM min^−1^, respectively (Table 3). The catalytic efficiency of CPB6-CoAT for caproyl-CoA was 3.8 times (41.08 vs 10.82 mM^−1^min^−1^) higher than that for butyryl-CoA, consistent with our previous result showing that the CCoAT activity is predominantly higher than the BCoAT activity (21). The *K*_m_ of CPB6-CoAT for caproyl-CoA was significantly lower than that for butyryl-CoA (358.8 vs 536.9 µM), exhibiting the higher affinity of this enzyme for caproyl-CoA relative to butyryl-CoA. These results also partly explained why caproate instead of butyrate is always the predominant product in the fermentation broth of strain CPB6 (22,23). BEY8-CoAT had only BCoAT activity, with *K*_m_, *k*cat, *k*_cat_/*K*_m_ and *V*_m_ values of 369.5 µM, 13.91 min^−1^, 37.65 mM^−1^min^−1^ and 27.81 μM min^−1^, respectively, and there was no detectable CCoAT activity (Tables 2 and 3), supporting our previous results showing that strain BEY8 produces only butyric acid as the predominant product (24). Similar to the results of Lee et al., the CoAT from *C. tyrobutyricum* only catalyzes the conversion of butyryl-CoA to butyrate and is not responsible for chain elongation of larger or higher-carbon-numbered (>C5) fatty acids (29).

**Table 3.** Kinetic parameters for the CoATs.

| Enzyme | Butyryl-CoA |  |  |  | Caproyl-CoA |  |  |  | Reference |
| --- | --- | --- | --- | --- | --- | --- | --- | --- | --- |
| | $K_m$<br>($\mu\text{M}$) | $k_{\text{cat}}$<br>($\text{min}^{-1}$) | $k_{\text{cat}}/K_m$<br>($\text{mM}^{-1}\text{min}^{-1}$) | $V_m$<br>($\mu\text{M min}^{-1}$) | $K_m$<br>($\mu\text{M}$) | $k_{\text{cat}}$<br>($\text{min}^{-1}$) | $k_{\text{cat}}/K_m$<br>($\text{mM}^{-1}\text{min}^{-1}$) | $V_m$<br>($\mu\text{M min}^{-1}$) | |
| BEY8-CoAT | 369.5 | 13.91 | 37.65 | 27.81 | ND<sup>c</sup> | ND<sup>c</sup> | ND<sup>c</sup> | ND<sup>c</sup> | This study |
| CPB6-CoAT | 536.9 | 5.810 | 10.82 | 11.62 | 358.8 | 14.74 | 41.08 | 29.51 | This study |
| CPB6-CoAT-D346H-mutant | 747.4 | 1.296 | 1.734 | 2.592 | 747.8 | 4.235 | 5.663 | 8.469 | This study |
| CPB6-CoAT-A351P-mutant | 622.7 | 2.987 | 4.797 | 5.975 | 532.1 | 6.784 | 12.75 | 13.57 | This study |
| PGN_0725<sup>a</sup> | 520.0 | 9.33 | 17.95 | 71.51 | NR<sup>c</sup> | NR<sup>c</sup> | NR<sup>c</sup> | NR<sup>c</sup> | Sato, Yoshida et al.<br>2016 (16) |
| CoA transferase<sup>b</sup> | 21.01 | NR<sup>c</sup> | NR<sup>c</sup> | NR<sup>c</sup> | NR<sup>c</sup> | NR<sup>c</sup> | NR<sup>c</sup> | NR<sup>c</sup> | Papoutsakis. 1989 (27) |

<sup>a</sup> Butyryl-CoA:acetate CoA transferase from *Porphyromonas gingivalis*.
<sup>b</sup> Butyryl-CoA:acetate CoA transferase from *Clostridium acetobutylicum* ATCC 824.
<sup>c</sup> ND is defined as not determined; NR, not reported.

The *K*_m_ of BEY8-CoAT for butyryl-CoA (369.5 µM) was obviously greater than that of CPB6-CoAT (536.9 µM), indicating that BEY8-CoAT had higher enzymatic affinity for butyryl-CoA than CPB6-CoAT. Similarly, CoAT (PGN_0725) from *Porphyromonas gingivalis* (15,16) and CoAT from *C. acetobutylicum* ATCC 824 (27) both catalyze the conversion of butyryl-CoA to butyrate, with *K*_m_ values of 520 µM and 21.01 µM, respectively. This indicated that the BCoAT generally had higher affinity and catalytic activity for butyryl-CoA than the CCoAT, while no BCoAT from butyric acid-bacteria displayed affinity and catalytic activity for caproyl-CoA. These results suggested that the BCoAT is only involved in chain elongation of C2 to C4, not in that of C4 to C6 or C8.

### Phylogenetics of the whole genome and multiple amino acid sequence alignment

The whole-genome phylogenetic tree was constructed based on 119 single-copy genes (including CoATs) that were common among 29 strains (Fig. 2). These strains have a wide range of butyrate metabolic pathways (29), for example, *Roseburia* sp., *Faecalibacterium prausnitzii*, and *Coprococcus* sp. from the human gut exhibit BCoAT activity values of 38.95, 18.64 and 118.39 U/mg of protein (crude extracts), respectively (13). The two species closest to strain CPB6 were *Pygmaiobacter massiliensis* (32) and *F. prausnitzii* (33), which are also butyric acid-producing bacteria in human feces. Interestingly, the species closest to *C. tyrobutyricum* BEY8 was *C. kluyveri*, which is a well-known caproic acid-producing bacterium. This close relationship may be because they belong to the same genus, *Clostridium*.

**FIGURE 2.**
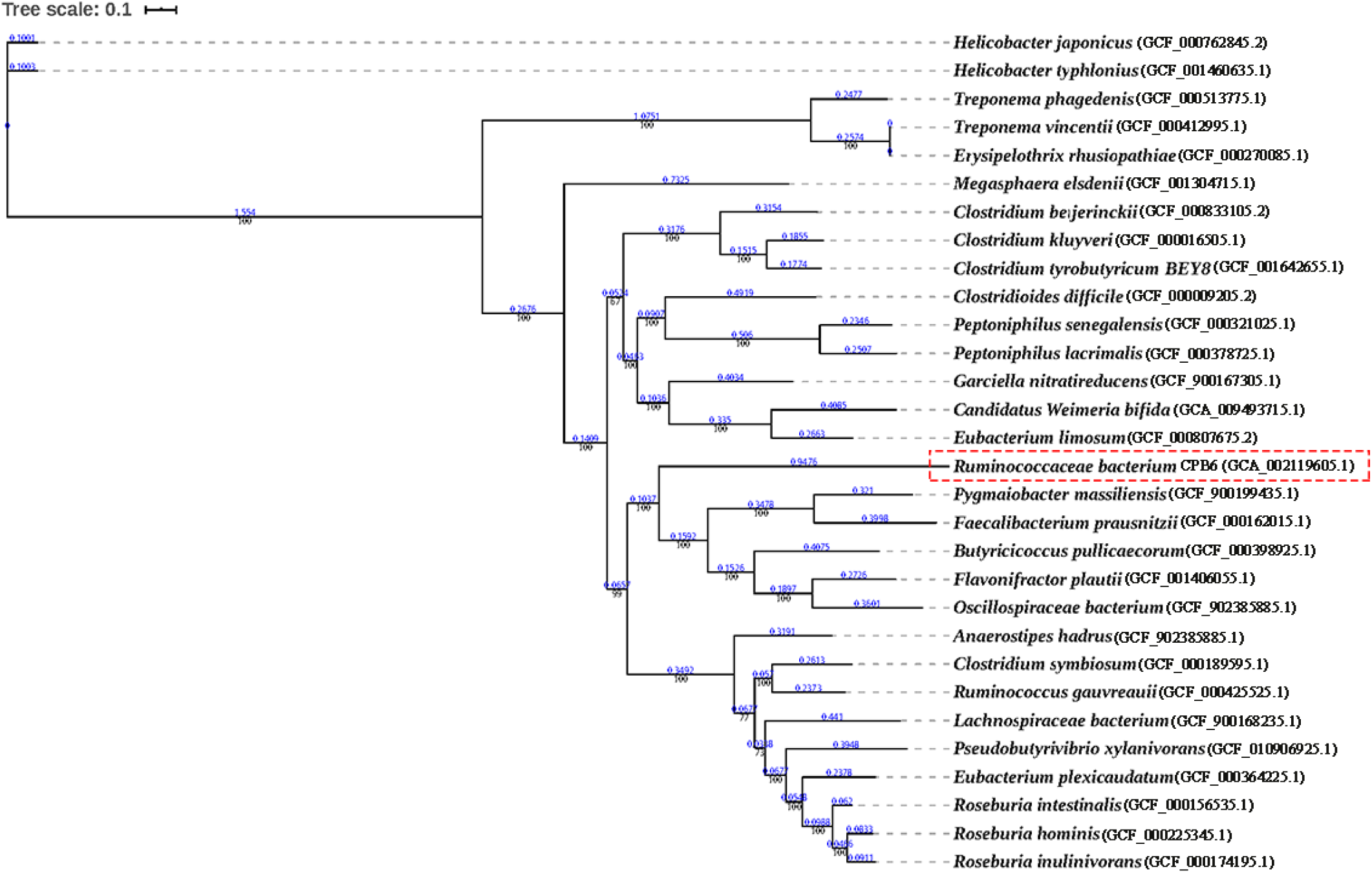
Phylogenetic tree of the whole genomes of 29 strains containing the CoA-transferase. Numbers at the nodes indicate the levels of bootstrap values. The scale bar for the tree represents a distance of 0.1 substitutions per site.

Based on the alignment results generated from CoAT protein sequences from six different species, 14 amino acids (GXGGQXDFXXGAXX, position 340-353) of the CoATs in all the microbes were highly conserved (except in *M. elsdenii*), and their secondary structures consisted of 17 α-helices and 21 β-sheets (Fig. 3). The sequence similarities between CPB6-CoAT and the analyzed CoATs were as follows: *C. kluyveri* (37.67%), *M. elsdenii* (10.27%), *C. tyrobutyricum* BEY8 (38.04%), *Lachnospiraceae* bacterium (60.59%), and *Anaerostipes hadrus* (58.52%). Among the six bacteria, strain CPB6, *C. kluyveri*, and *M. elsdenii* are caproic acid-producing bacteria, while *C. tyrobutyricum* BEY8, *Lachnospiraceae* bacterium, and *A. hadrus* are butyric acid-producing bacteria. The alignment results showed that CPB6-CoAT shared lower similarity (10.27-37.67%) with the CoATs of *C. kluyveri* and *M. elsdenii* but higher similarity (58.52-60.59%) with the CoATs of *Lachnospiraceae* bacterium and *A. hadrus*. This may be because strain CPB6 belongs to the family *Ruminococcaceae*, which is closer to *Lachnospiraceae* and *Anaerostipes* at the taxonomic phylogeny level than *Megasphaera* and *Clostridium*.

**FIGURE 3.**
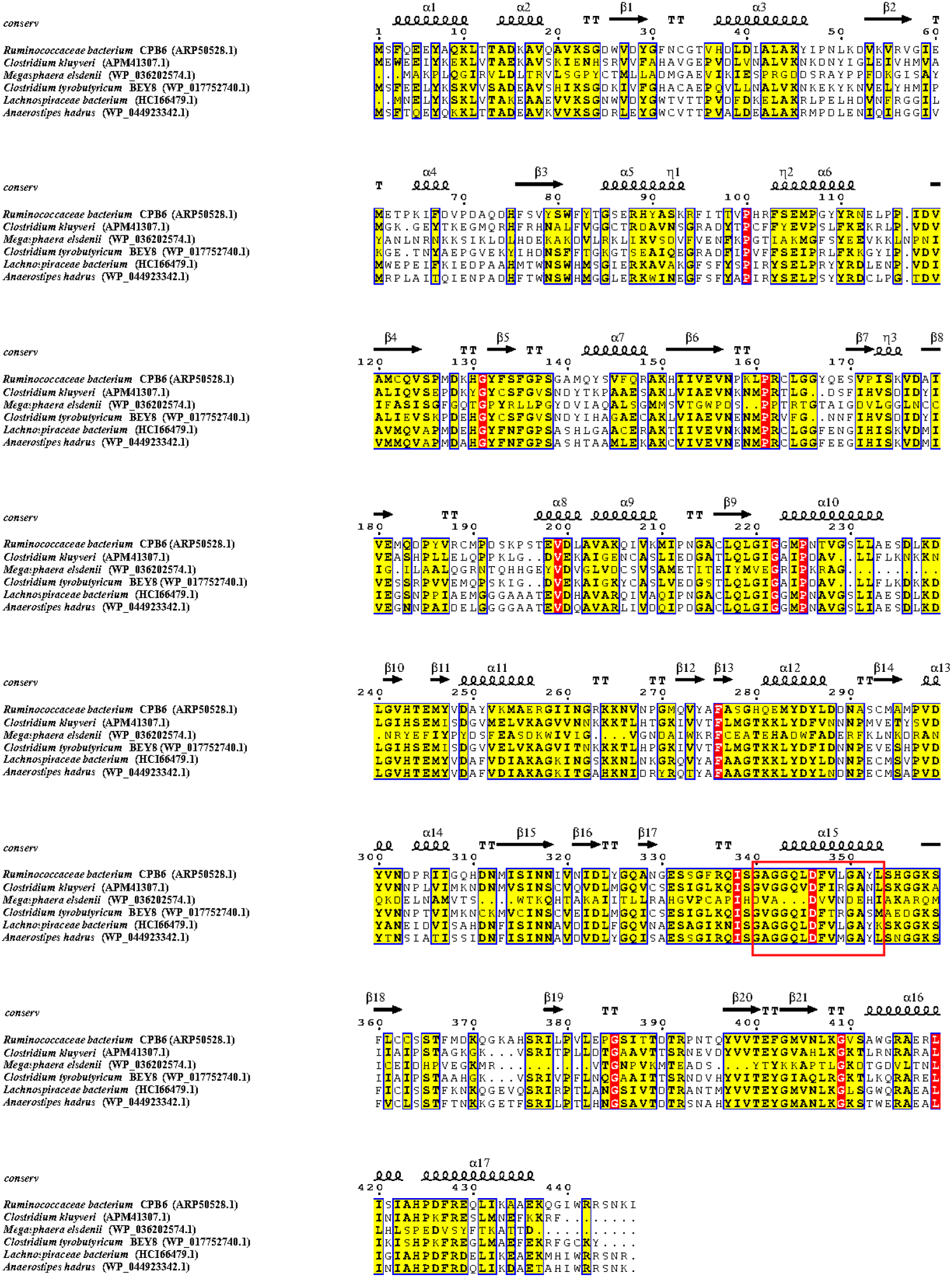
Multiple amino acid sequence alignment for CoATs. There were 17 α-helices and 21 β-pleated sheets, which are represented with symbols. Nonconserved, 60% conserved, and 100% conserved residues are marked with white, yellow, and red font, respectively. Conserved motifs are boxed with a red frame.

### Prediction and comparison of the three-dimensional (3D) structure and active site

As shown in Fig. 4 *A*, the three-dimensional (3D) structure of CPB6-CoAT has one subunit which may consist of two main domains resulting in a characteristic two -domain fold in a homo-tetrameric structure. A comparison of the 3D structures of the six CoAT proteins (Fig. 4) showed that these CoATs shared similar conformations of the structural elements (α-helices and β-strands) with slight structural modifications in the loop regions and active centers, with the exception of the CoAT from *M. elsdenii* (Fig. 4*C*). The 3D structure of the *M. elsdenii* protein was obviously different from that of other CoATs, and the divergences were located not only in the structural elements of α-helices and β-strands but also in the loops. This may be attributed to the distant genetic relationship between *M. elsdenii* and the other five bacteria. Although *M. elsdenii* produces caproic acid via acetyl-CoA and succinate (34), the functions of the CoATs may differ between strain CPB6 and *M. elsdenii*. The 3D structures of CoATs among *C. tyrobutyricum* BEY8, *Lachnospiraceae* bacterium, and *A. hadrus* shared almost the same conformation of α-helices and β-strands except for some slight variation in the loops (Fig. 4*D–F*). The protein structure and active center structures between CPB6-CoAT and BEY8-CoAT were further compared, as shown in Fig. S4, and both showed similar 3D structures except for the location and structure of the active center. The predicted active sites of the six CoATs are shown in Table 4. The predicted active center of the CPB6 -CoAT protein was located between amino acids 342 and 353 (GGQLDFVLGAYL) while the active center of the BEY8-CoAT protein (GGQIDFTRGASM) was located from amino acids 335 to 346, and both active site peptides contain a phenylalanine and tyrosine (Fig. S4).

**FIGURE 4.**
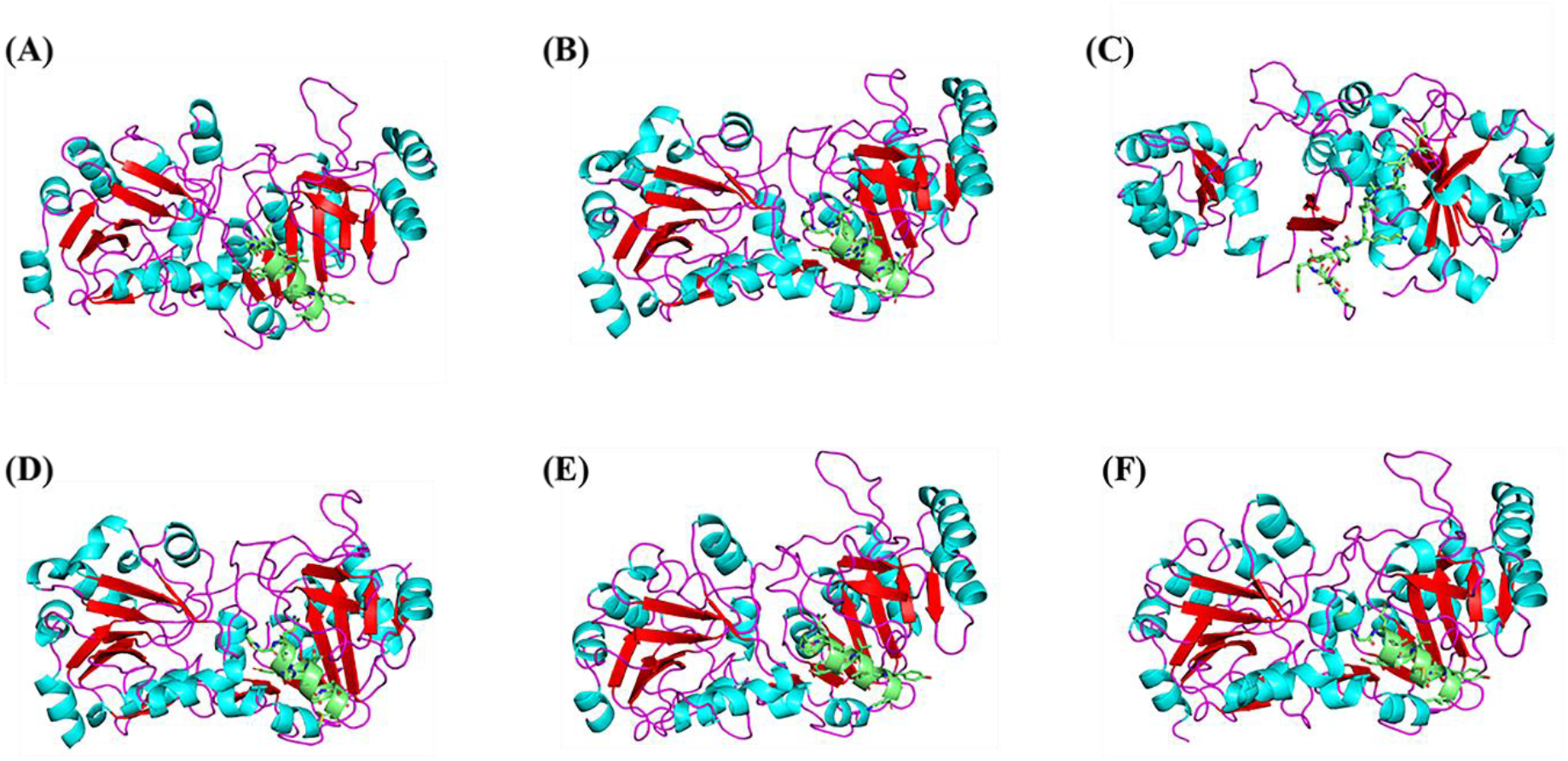
Predicted 3D structures of representative CoAT proteins. 3D structures of CoATs from *Ruminococcaceae* bacterium CPB6 **(A)**, *Clostridium kluyveri* **(B)**, *Megasphaera elsdenii* **(C)**, *Clostridium tyrobutyricum* BEY8 **(D)**, *Lachnospiraceae* bacterium **(E)**, and *Anaerostipes hadrus* **(F)**. Helices of the catalytic domains, β-pleated sheets, loop regions, and active centers are colored sky blue, red, purple, and green, respectively.

**Table 4.** Prediction of the active sites of CoATs in different strains.

| Strain | Location of active site | Sequence of active site |
| --- | --- | --- |
| *Ruminococcaceae* bacterium CPB6 (ARP50528.1) | 342-353 | GGQLDFVLGAYL |
| *Clostridium kluyveri* (APM41307.1) | 335-346 | GGQVDFIRGANL |
| *Megasphaera elsdenii* (WP_036202574.1) | 356-367 | ADSYTYKKAPTL |
| *Clostridium tyrobutyricum* BEY8 (WP_017752740.1) | 335-346 | GGQIDFTRGASM |
| *Lachnospiraceae* bacterium (HCI66479.1) | 340-351 | GGQLDFVLGAYK |
| *Anaerostipes hadrus* (WP_044923342.1) | 342-353 | GGQLDFVMGAYL |

The structure of proteins plays an important role in their functional properties and catalytic efficiency (35); for example, succinyl CoA:3-ketoate CoA transferase from pig heart (36) and 4-hydroxybutyrate CoA-transferase from *Clostridium aminobutyricum* (37) showed unexpected changes in protein modification and specific activity when their crystal structures changed. In this study, a comparison of the 3D and active center structures showed the similarities and differences between CPB6-CoAT and other CoATs (Fig. 4 and S4), which may have affected the enzyme catalytic function and activity. On the basis of these results, it is of great significance to study the functional differences caused by the structural changes in CoATs. The exact structure and function of the active center of the CPB6-CoAT protein remains to be determined through subsequent comprehensive experiments and analysis.

### Site-directed mutagenesis

Site-directed mutagenesis was used to verify the active sites of the protein (38). According to the predicted active center of CPB6-CoAT (GGQLDFVLGAYL, 342-353 aa), site-directed mutagenesis targeting sites Asp346 and Ala351 was carried out to identify the effects of the two residues on the catalytic activity of CPB6-CoAT. Specifically, Asp346 was replaced by His and Ala351 was replaced by Pro via site-directed mutagenesis. The nucleotide substitutions were confirmed by Sanger sequencing of the DNA (Fig. S5). Enzyme assays showed that compared to wild-type CPB6-CoAT, the Asp346 substitution led to an approximately 76% loss of BCoAT activity and 72% loss of CCoAT activity, while the Ala351 substitution resulted in an almost 50% loss of BCoAT activity and 55% loss of CCoAT activity (Fig. 5). Moreover, as shown in Table 3, the *k*cat/*K*_m_ values for butyryl-CoA and caproyl-CoA of the D346H mutant (1.734 and 5.663 mM^−1^min^−1^) and A351P mutant (4.797 and 12.75 mM^−1^min^−1^) were all lower than those of wild-type CPB6-CoAT (10.82 and 41.08 mM^−1^min^−1^). This result indicated that the Asp346 and Ala351 residues play vital roles in the active center of CPB6-CoAT.

**FIGURE 5.**
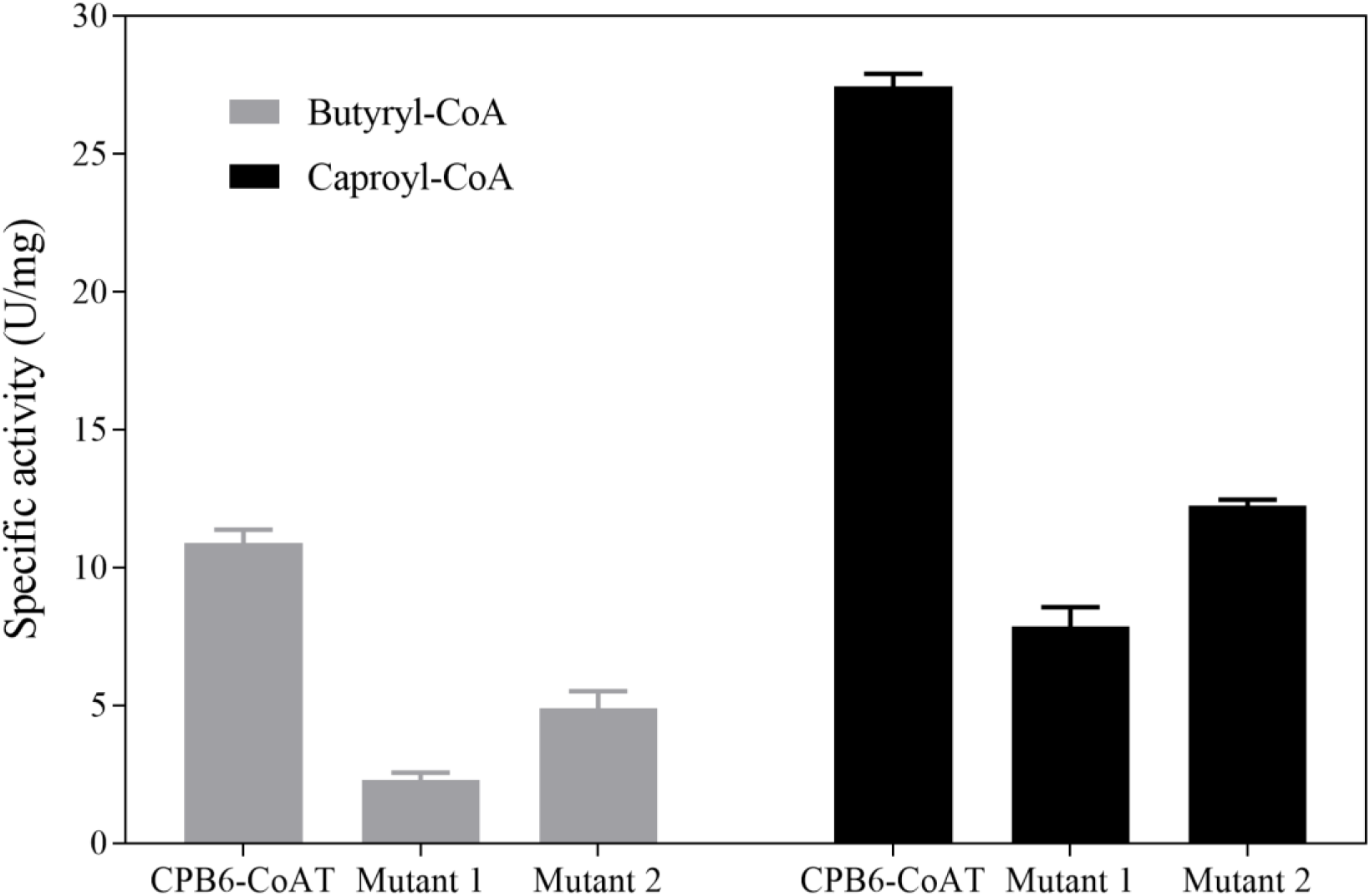
Comparison of CoA-transferase activities. (Mutant 1, D346H-mutant; Mutant 2, A351P-mutant). The specific activity with butyryl-CoA as the substrate is labeled in light gray, and the specific activity with caproyl-CoA as the substrate is marked in dark gray. The values represent the means ± SDs of three independent experiments.

Notably, the exchange of Asp, an acidic amino acid, for His led to loss of a carboxy group and the introduction of two amidogens. Similarly, the replacement of Ala with Pro led to loss of an amidogen and the introduction of a carboxy group. Ala lacked a bulky side chain and therefore would likely not have any steric and electrostatic effects, and this change would not destroy the conformation of the main chain (39). Differences in structures and properties among the sequences may be the reason for the differences in CoAT activity (40). These results demonstrated that the conserved motif (GGQLDFVLGAYL, 342-353 aa) of CPB6-CoAT is directly linked to enzymatic activity. In conclusion, the conserved motif was the catalytic center of CPB6-CoAT, within which the Asp346 and Ala351 residues were essential for CoAT activity. These findings provide significant information for further detailed research on the structures and functions of CPB6-CoAT.

#### Concluding remarks

In our present study, two CoA-transferase genes were identified by cloning and expression in *E. coli* BL21 (DE3) with the plasmid pET28a. CPB6-CoAT showed higher activity for caproyl-CoA than for butyryl-CoA, while BEY8-CoAT only had activity for butyryl-CoA. This result indicated that CPB6-CoAT is responsible for the final step of caproic acid production. The bioinformatics analysis revealed differences between CPB6-CoAT and other CoATs. Moreover, site-directed mutagenesis analysis demonstrated that the conserved motif (GGQLDFVLGAYL, 342-353 aa) was probably the active center of CPB6-CoAT, within which sites Asp346 and Ala351 residues were identified as critical residues that affect the enzymatic activity of CPB6-CoAT. These results confirmed the existence of a CCoAT involved in the production of caproic acid, and the enzyme is apparently different from the BCoAT responsible for the production of butyric acid. This study facilitates our understanding of the metabolism for chain elongation via the reverse β-oxidation pathway. However, the detailed CCoAT structure and its function in MCFA biosynthesis require further study through crystallization of proteins and X-ray crystal structure analysis.

### Experimental Procedures

#### Strain growth conditions

*E. coli* DH5α (TsingKe, Chengdu, China) and *E. coli* BL21 (DE3) (Transgene, Beijing, China) were cultured in Luria broth (LB) medium supplemented with 50 µg/ml kanamycin (Sangon Biotech, Shanghai, China) at 37 °C. *Ruminococcaceae* bacterium CPB6 was grown anaerobically at 37 °C in modified reinforced Clostridium medium (Binder, Qingdao, China) (20). *C. tyrobutyricum* BEY8 was grown anaerobically at 37 °C in TGY medium (30 g/L tryptone, 20 g/L glucose, 10 g/L yeast extract, and 1 g/L L-cysteine hydrochloric acid; pH 7.0).

#### Gene cloning and plasmid construction

The CoAT genes were amplified from the genomic DNA of *Ruminococcaceae* bacterium CPB6 (23) or *C. tyrobutyricum* BEY8 (24) through PCR using the primers listed in Table 1. During amplification, the following conditions were used: initial denaturation (5 min at 98 °C), followed by 30 cycles of denaturation (10 s at 98 °C), annealing (30 s at 52 °C), and elongation (1 min at 72 °C) and a final extension (5 min at 72 °C). The PCR products were verified by agarose electrophoresis, recovered using a PCR purification kit (Fuji, Chengdu, China), and seamlessly inserted into the plasmid pET28a double digested with *Not* I and *Sal* I (Thermo, Waltham, USA) to construct the recombinant plasmid by using a seamless cloning kit (Biomed, Beijing, China). The recombinant plasmids were verified by Sanger sequencing and then transformed into *E. coli* BL21 (DE3) cells.

#### Expression and purification of the CoA-transferases

The recombinant plasmids pET28-CoAT-CPB6 (pET28-CCoAT) and pET28-CoAT-BEY8 (pET28-BCoAT) were transformed into *E. coli* BL21 (DE3). The transformed cells were cultured in LB medium containing 50 µg/ml kanamycin at 37 °C until the OD_600_ reached 0.5 and then further cultured at 22 °C for 12 h with 0.4 mM IPTG. The cultured cells were harvested by centrifugation (8,000 × *g*, 10 min) at 4 °C, and the cell pellet was resuspended in 50 mM potassium phosphate (pH 8.0). The cells were then disrupted by an ultrasonicator (Huxi, Shanghai, China) for 30 min (200 W, 4 s, interval 6 s) and centrifuged at 8,000 × *g* for 30 min to remove the insoluble material. Then, the enzyme was purified with Ni-NTA Sepharose (Genscript, Nanjing, China) and eluted with 50 mM sodium phosphate (pH 8.0) containing 300 mM NaCl and 250 mM imidazole. Finally, the purity and MW of the enzyme were assessed using SDS-PAGE analysis. Moreover, the enzyme was analyzed by western blotting with anti-6×His rabbit polyclonal antibody (Sangon Biotech, Shanghai, China). The protein concentrations were determined using a BCA protein assay kit (Solarbio, Beijing, China).

#### Enzymatic characterization

CoAT activity in crude enzyme extracts and of purified recombinant proteins was measured by determining the concentration of acetyl-CoA, a reaction byproduct, with the citrate synthase assay as described in previous studies with minor modifications (25,41). In brief, the reaction was initiated by the addition of enzyme (up to 20 ng/ml) and was performed in a total volume of 1.0 ml at 25 °C: 100 mM potassium phosphate buffer (pH 7.0), 200 mM sodium acetate, 1.0 mM 5,5’-dithiobis (2-nitrobenzoate), 1.0 mM oxaloacetate, 8.4 *n*kat citrate synthase (Sigma, St. Louis, USA), 0.5 mM CoA derivatives (Sigma, St. Louis, USA). The released CoA, corresponding to the residual amount of acetyl-CoA, was detected by measuring the absorbance at 412 nm. One unit of activity is defined as the amount of enzyme which converts 1 µmol of acetyl-CoA per min under these conditions.

The kinetic parameters of the recombinant protein were also calculated by using the coupled spectrophotometric enzyme assay through citrate synthesis (16). The reaction mixture was the same as that mentioned above, and the concentrations of butyryl-CoA or caproyl-CoA were varied from 0.5 to 5 mM. The kinetic parameters were computed by using the Lineweaver–Burk transformation of the Michaelis-Menten equation, in which velocity is a function of the substrate (42,43). The catalytic constant (*k*_cat_) was defined as the number of CoAT molecules formed by one molecule of enzyme in a single second. All measurements were performed in triplicate for biological replications.

#### Bioinformatics

Sequence alignment was performed using ESPript (44). Representative CoAT (8,45-47) sequences were downloaded from the NCBI database. MEGA-X software was used to perform sequence alignment and construct the phylogenetic tree (48). The active sites of CoATs were predicted by the online tool ScanProsite, and the three-dimensional CoAT structure was simulated by NCBI-CDD to search for templates in SWISS-MODEL and was embellished and labeled by PyMOL 2.3.3 software (49,50). The phylogenetic relationships of CoATs from different species were obtained by using OrthoFinder, the amino acid sequences were downloaded from the NCBI website (version 2.2.7) (51), and MUSCLE (v3.8.31) was used to calibrate the 119 shared single-copy genes (52). The phylogenomic tree was derived from a supermatrix comprising these shared single-copy genes with 41,213 unambiguously aligned amino acids using the maximum likelihood (43) method in RAxML (v8.2.10) (53) under the PROTGAMMAAUTO model, with 100 bootstrap replicates.

#### Site-directed mutagenesis

The point mutation vectors were constructed with the Fast Mutagenesis System (Transgene, Beijing, China). The QuikChange PCR method using pfu DNA polymerase was performed to generate the D346H mutant and A351P mutant. The recombinant plasmid (pET28a-CoAT-CPB6) was used as template DNA, and complementary mutagenic oligonucleotides as primers are shown in Table 1. After PCR amplification, the mixture was digested with restriction enzymes using D*pn*I to remove methylated template DNA and then sequenced (TsingKe, Chengdu, China) to verify site mutagenesis before being transformed into *E. coli* BL21 (DE3) (Transgene, Beijing, China). After purification, the enzymatic activities for butyryl-CoA and caproyl-CoA were measured following the method described above for the wild type.

## Supplemental material

Supplemental material is available online only.

## Acknowledgements

This work was supported by the Natural Science Foundation of China (31770090), the Open-foundation project of CAS Key Laboratory of Environmental and Applied Microbiology (KLCAS-2017-01). Additionally, we would like to thank Dr. Su Dan for her advice and assistance in this study.

## Conflict of interest

The authors are aware of no conflict of interest.

